# Expression of immune checkpoint VISTA represents a differentiated state of cancer cells and plays a role in regulating actin cytoskeleton

**DOI:** 10.64898/2026.08.24.746888

**Authors:** Chen Wang, Yang Liu, Jing Li, Ying Cao

## Abstract

Immune checkpoint blockade has revolutionized cancer therapy, but the therapeutic efficacy is limited. Clinical trials on blockade of newly identified immune checkpoints didn’t show promising result, suggesting that it might be insufficient to understand the function of immune checkpoints in cancer merely in the context of immunity. Here, we found mutually exclusive expression patterns of the immune checkpoint VISTA (or VSIR) and the neural stemness factor SETDB1, an oncoprotein that promotes immunoevasion, in xenograft tumors, suggesting that cells with high VISTA expression represents a differentiated, and hence, less or non-malignant state in tumor. Non-neural differentiation factors HHEX, MYOD1 and PPARG promote, whereas oncoproteins KRAS (and the mutant KRAS(G12D)) and SOX2, both being embryonic neural factors, repress VISTA expression. This tendency can be inferred from the finding that neural stemness is the core property of cancer cell. Manipulated expression of VISTA in cancer cells generated no significant effect on cell tumorigenicity and differentiation state, but led to change in cell morphology and actin cytoskeleton. Mechanistically, VISTA regulates a key cytoskeleton regulator, WASF2, leading to the change in cell morphology, which might interfere with signal transduction of immune response. The results suggest that 1) high expression of a protein in tumor might represent a less or non-malignant state, targeting of which would leave malignant cells intact, and consequently, leading to weak or even no therapeutic efficacy, a key factor worth considering for target selection; 2) immune checkpoints might play other roles in cells that interfere with regulation of anti-tumor immunity.

## Introduction

Cancer is the most extensively investigated disease so far. Despite the fact that numerous molecular mechanisms underlying cancer have been elucidated, the basic property of cancer cells, cancer and cancer complexity were still poorly understood. Our previous studies demonstrated that most cancer promoting genes are embryonic neural/neural stemness genes and suppressor genes are mainly not, and the core property of cancer (tumorigenic) cells is neural stemness. Neural stemness represents the basal cellular property that confers cancer cells with pluripotency and tumorigenicity, and unifies cancer cell phenotypic traits, including stemness, migration, immune evasion, etc. (Zhang et al., 2017; Cao, 2017; Lei et al., 2019; Xu et al., 2021; Chen et al., 2021; Cao, 2022; Zhang et al., 2022; Cao, 2023; Cao, 2026). This means that cancer cells can differentiate into different types of cells, including neuronal cells and cells in non-neural lineages, similar to pluripotent embryonic cells, and differentiated cancer cells loses their neural stemness and malignant features, and acquirement of immunogenicity (Zhang et al., 2017; Xu et al., 2021; Yang et al., 2021; Cao, 2022; Zhang et al., 2022; Cao, 2023; Linde et al., 2023; Zimmermannova et al., 2023;Ascic et al., 2024; Gong et al., 2025; Cao, 2026). Besides cell-intrinsic cancer promoting genes, many other genes have also been identified to express in a wide spectrum of cancers and their expression is positively correlated with poor prognosis and poor response to treatment, for example, TUBB3 (Kanakkanthara and Miller, 2021) or ITGB6 (Brzozowska and Deshmukh, 2022). They are also considered as strong therapeutic targets for cancer treatment. Nevertheless, if considering that TUBB3 is a typical marker indicating differentiated neuronal cells and ITGB6 is muscle-specific during embryogenesis (Thisse and Thisse, 2004), then high level of expression of these genes should indicate a differentiated state and tumor cells with high level of expression of these genes should be of weaker or no tumorigenic potential. This raises a question what it means when a gene is found to express highly in cancer and correlate with poor prognosis and pathological grades, and whether this is an efficient criterion for target selection.

Immune checkpoint blockade removes inhibitory signals of T-cell activation, typically PD-1/PD-L1 and CTLA-4, thereby reactivating the immune system to recognize and destroy cancer cells. After the initial success of PD-1/PD-L1 and CTLA-4 blockade in cancer therapy, a series of novel immune checkpoints have been identified, including LAG-3, TIGIT, IDO1, TIM-3, and VISTA. They have been expected to improve the efficacy of PD-1/PD-L1or CTLA-4 blockade therapy because only a very limited number of cancer patients can benefit from the therapy, resistance also develops after therapy, T cell exhaustion occurs, or tumor microenvironment (TME) helps cancer cells evade from immune killing. Sometimes this therapy even causes hyperprogression of cancer (Kamada et al., 2019; de Miguel and Calvo, 2020; Morad et al., 2021; Aliazis et al., 2025). However, clinical trials with blockade of these novel immune checkpoints, IDO1, TIGIT, or LAG-3, have shown little or no benefit in decreasing tumor burden or shifting survival (Long et al., 2019; Yang and Chen, 2025; Mullard, 2026; Zhang et al., 2026). The costly lessons from these trials raise another question whether it is sufficient to understand the function of immune checkpoints merely in the context of immunity.

The immune checkpoint VISTA (also known as PD-1H, B7-H5, VSIR, or Dies-1) acts as a co-inhibitory molecule that suppresses T-cell activation, proliferation, and cytokine production. It functions bidirectionally: as a ligand on antigen-presenting cells /myeloid cells that engages receptors on T cells, and as a receptor on T cells themselves. In the tumor microenvironment, VISTA contributes to immune evasion by suppressing antitumor T-cell responses and supporting an immunosuppressive milieu. Under experimental settings, its blockade can enhance antitumor immunity (Wang et al., 2011; Lines et al., 2014; ElTanbouly et al., 2020; Zhang and Kim, 2024; Abooali et al., 2025). These features of VISTA in regulating anti-tumor immunity laid the foundation for clinical trials of VISTA blockade. VISTA expression shows heterogeneous patterns in different types of cancer. Significant increase in VISTA expression was observed in cholangiocarcinoma, glioblastoma, kidney renal clear cell carcinoma, acute myeloid leukemia, low-grade gliomas, and pancreatic ductal adenocarcinoma as compared to paired normal tissue, but lower expression relative to paired normal tissues was detected in diffuse large B-cell lymphoma and melanoma. The heterogeneity in expression opens the possibility to consider VISTA expression as a potential biomarker for efficacy (Martin et al., 2023). Besides VISTA expression in tumor cells and immune cells, it is also expressed in a wide range of normal tissues, including muscle (Martin et al., 2023), raising the possibility of its function beyond immunity. In the present research, we showed correlation of VISTA expression with differentiation state of cancer cells, and its potential role in regulating actin cytoskeleton, thereby regulating cell morphology.

## Results

### Expression of VISTA represents a differentiated cell state in tumors

Xenograft tumors were obtained by subcutaneous injection of colorectal cancer cell HCT116, leukemia cell K562, liver cancer cell SK-HEP1, lung cancer cell NCI-H460, melanoma cell A375, or pancreatic cancer cell BxPC3, into immunodeficient nude mice. Immunohistochemistry (IHC) was performed to detect expression of SETDB1, which represents stemness or undifferentiated state and the state of immunoevasion (Johnson et al., 2023; Kang, 2015; Zhang et al., 2022), and VISTA, in two immediate adjacent sections, so as to compare their distribution of expression in tumors. In tumors formed by HCT116 cells, it was obvious that SETDB1 was primarily detected in nuclei of densely arranged cells with larger nuclei but not in smaller cell nuclei, showing an undifferentiated state. In the adjacent section, VISTA showed a complementary expression pattern because it was detected only in loosely arranged cells with smaller nuclei, indicating a differentiated state (Fig. 1A). These exclusive patterns of expression of SETDB1 and VISTA in undifferentiated and differentiated cells were also detected in tumors formed by K562, SK-HEP1, NCI-H460, A375, and BxPC3 cells (Fig. 1A). We also detected the expression patterns of Setdb1 in tumors formed by mouse NE-4C neural stem cells, B16F10 melanoma cells and 4T1 breast cancer cells. It is clear that Setdb1 was detected in undifferentiated cells in tumors derived from NE-4C cells and Vista was in differentiated cells (Fig. 1B). Similar exclusive expression of Setdb1 and Vista was also obvious in tumors formed by B16F10 or 4T1 (Fig. 1B). These results showed a previously unrecognized expression pattern of the immune checkpoint VISTA in differentiated cancer cells only, suggesting that expression of VISTA represents a differentiated state in tumors. Expression of SETDB1 indicates an undifferentiated and malignant cell state, but expression of VISTA represents a differentiated and less or non-malignant state.

**Fig. 1.**
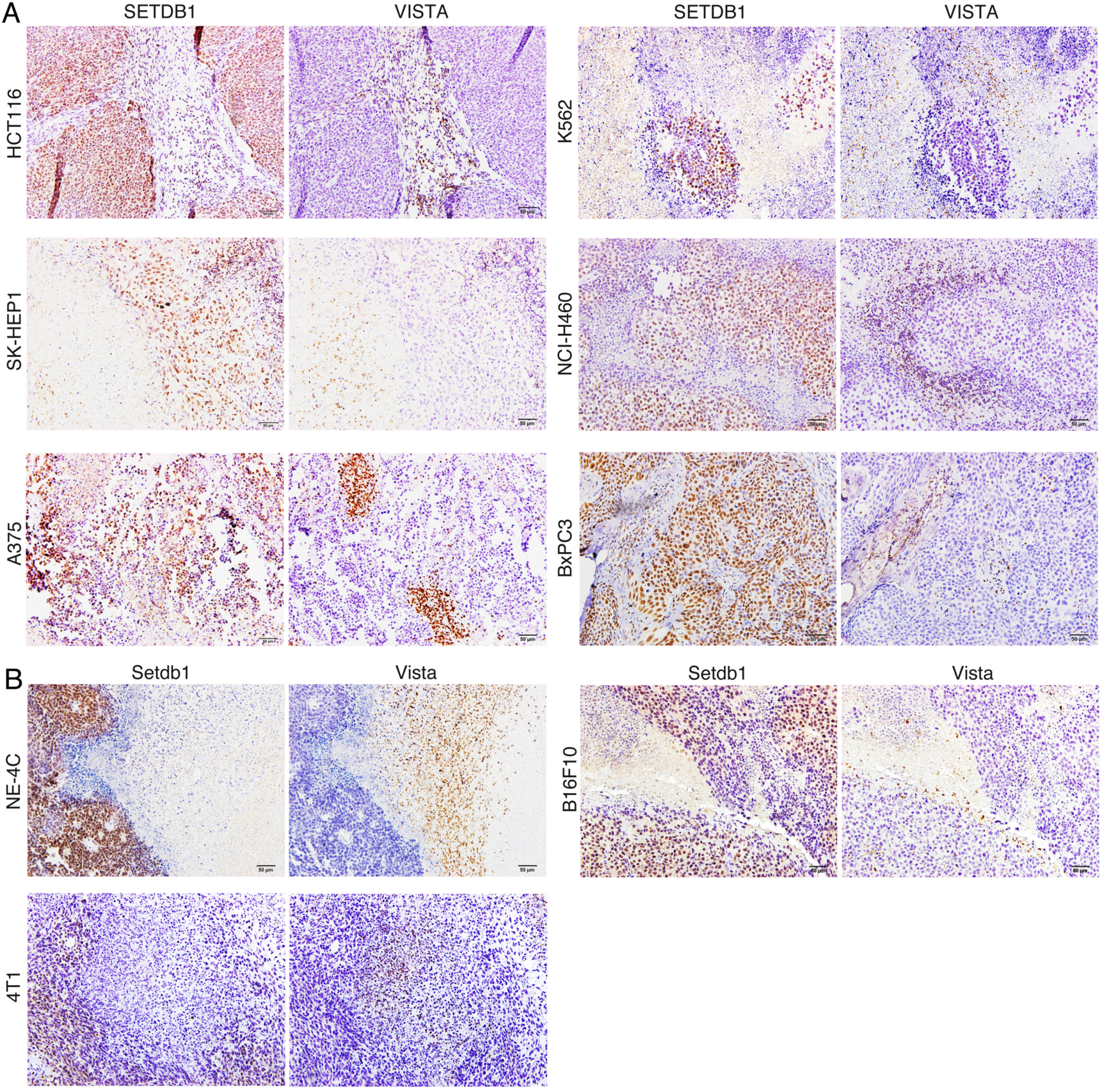
Mutually exclusive expression patterns of SETDB1/Setdb1 and VISTA/Vista in xenograft tumors. (A) IHC detection of SETDB1 and VISTA in two adjacent sections of tumors formed by six types of human cancer cells. (B) IHC detection of Setdb1 and Vista in two adjacent sections of tumors formed by neural stem cells NE-4C and mouse B16F10 and 4T1 cancer cells.

### Phenotypic change in cells in response to inhibition or overexpression of VISTA

Next, we examined whether VISTA plays certain roles in regulating cancer cell differentiation. A short-hairpin RNA against VISTA (shVISTA) was used to knock down VISTA in cancer cells via lentiviral transfection. Interestingly, A375 cells showed strong phenotypic change in response to knockdown of VISTA. Compared with control, cells with VISTA knockdown exhibited a spherical morphology. Immunoblotting revealed that shVISTA could inhibit VISTA expression efficiently. However, VISTA knockdown could only generate weak effect on expression of the cell proliferation marker PCNA, neural stemness markers MSI1 and SETDB1 (Fig. 2A). High expression of these proteins indicates high malignancy of cancer cells. Therefore, change in expression of these factors after VISTA knockdown suggested a weak change in malignancy. In HCT116 cells, knockdown of VISTA led to formation of small clusters, and reduced expression of oncoproteins SETDB1, EZH2, but slightly enhanced expression of MSI1 (Fig. 2A). Weak changes in the morphology of SK-HEP1 and NCI-H460 cells were observed after knockdown of VISTA. Accordingly, expression of PCNA, MSI1 and oncoproteins SETDB1, EZH2, or MYC was not changed significantly (Fig. 2A).

**Fig. 2.**
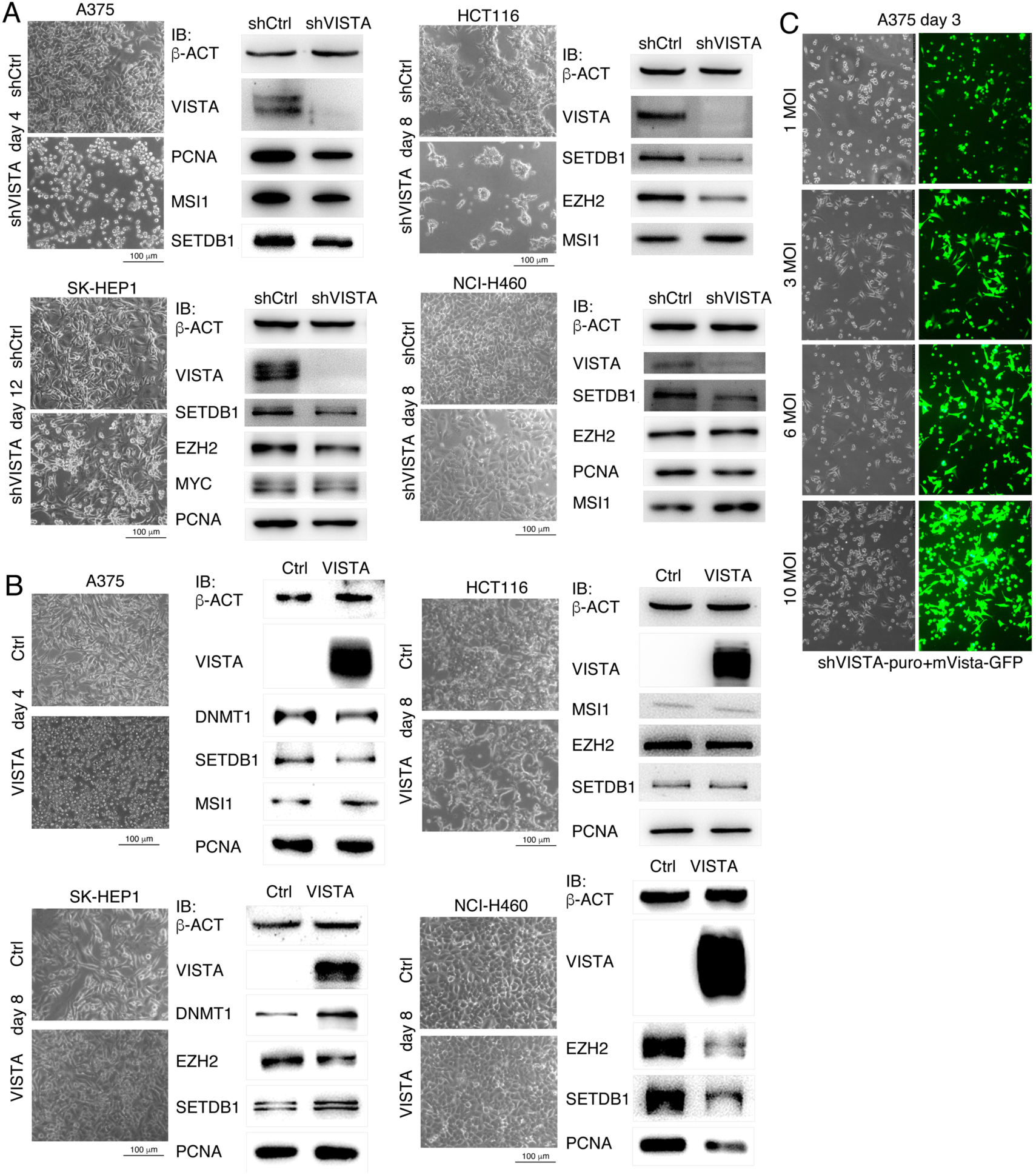
Manipulated expression of VISTA in cancer cells caused cell morphological change. (A) Phenotypic and protein expression changes in cancer cells with knockdown of VISTA. In each experiment, control (shCtrl) and knockdown (shVISTA) cells were collected and subjected to immunoblotting (IB). β-ACT was used as a loading control. (B) Phenotypic and protein expression changes in cancer cells in response to overexpression of VISTA. In each experiment, control cells (Ctrl) and cells with VISTA overexpression (VISTA) were collected and subjected to IB. β-ACT was used as a loading control. (C) Rescue of the morphological alteration in A375 cells resulting from VISTA knockdown by overexpression of mouse Vista (mVista) via transfection of virus at different MOIs as indicated.

To examine VISTA function in cells further, overexpression of VISTA in cancer cells was also performed via lentiviral transfection. Intriguingly, overexpression of VISTA caused a significant change in A375 cells similar to VISTA knockdown because the cells also assumed a spherical morphology. However, expression of oncoproteins and markers indicating cancer cell malignancy was not altered significantly in A375 cells after overexpression (Fig. 2B). We didn’t observe significant change in cell morphology and protein expression in HCT116, SK-HEP1, and NCI-H460 after VISTA overexpression. Knockdown and overexpression assays suggest that VISTA might play a role in regulating cell morphology.

Both knockdown and overexpression of VISTA caused a similar morphological change in A375 cells. To exclude this was an artifact, we examined whether mouse Vista (mVista) was able to rescue the phenotype of VISTA knockdown (shVISTA) in A375 cells. When the ratio of lentivirus for mVista to lentivirus for shVISTA was at 1 MOI (multiplicity of infection), most cells showed spherical morphology, showing no significant rescue effect. When the ratio was increased to 3 MOI, significant rescue effect was observed because a majority of cells did not show spherical morphology anymore. But when the ratio of two types of lentivirus was increased to 6 or 10 MOI, rescue effect decreased because more cells became spherical again (Fig. 2C). The rescue experiment demonstrates that morphological change in cells after VISTA knockdown or overexpression is specific.

Next we examined whether VISTA expression is required for cancer cell tumorigenicity. We observed that overexpression of VISTA in A375 or NCI-H460 cells did not change their tumorigenicity because no significant difference in xenograft tumor formation was observed, as compared with control cells (Fig. 3A). VISTA was identified as a key immune checkpoint in cancer immunity. However, we also did not observe that overexpression of Vista in mouse melanoma cell B16F10 or colorectal cancer cell CT26 led to a significant change in tumor formation in syngeneic mouse models (Fig. 3A). Vice versa, no significant change in tumor formation was observed in A375 and HCT116 cells with knockdown of VISTA (Fig. 3B). We did not test the effect of Vista knockdown on tumorigenicity of B16F10 and CT26 cells because Vista expression in these cells was barely detectable. The results suggest that VISTA is not a major factor regulating cancer cell tumorigenicity.

**Fig. 3.**
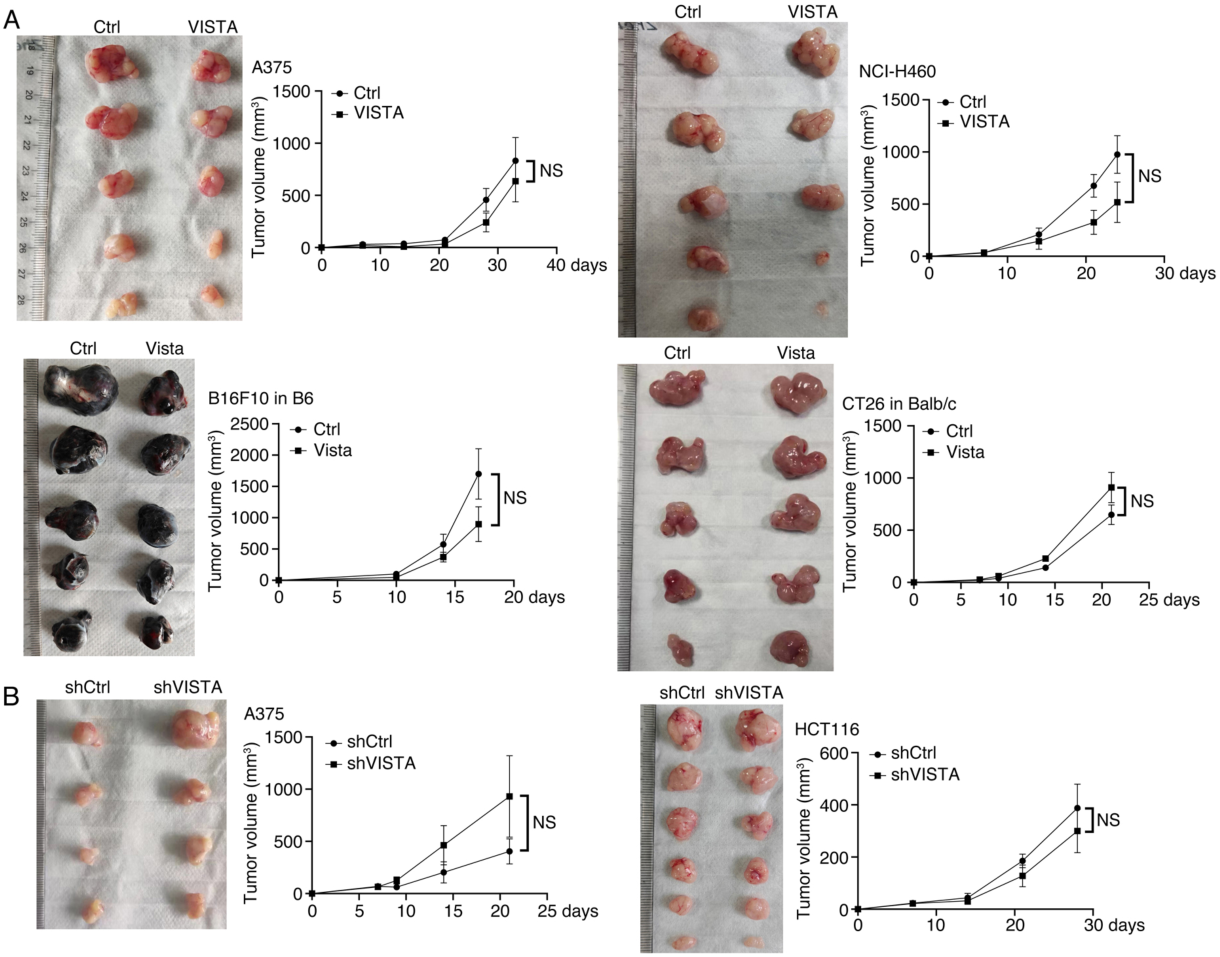
Effect of manipulated VISTA/Vista on cancer cell tumorigenicity. (A) Tumor formation by transplanting control cells (Ctrl) or cells with overexpression of VISTA/Vista into immunodeficient nude mice or syngeneic B6 or Balb/c mice. (B) Tumor formation by transplanting control (shCtrl) or knockdown (shVISTA) cells into immunodeficient nude mice. In (A) and (B), significance of difference in tumor volumes between two groups of mice was calculated using two-way ANOVA-Bonferroni/Dunn test. Data are represented as mean ± SEM. NS: Not significant.

### Differential regulation of VISTA by differentiation and dedifferentiation factors

Expression patterns of VISTA in xenograft demonstrate that VISTA is primarily expressed in differentiated cells. It can be deduced that VISTA expression could be stimulated by factors promoting differentiation but inhibited by factors that inhibit differentiation. Because the core property of cancer cells is neural stemness, which confers cancer cells with pluripotent differentiation potential, cancer cells can be driven to differentiate into a more mature state by non-neural differentiation factors (Yang et al., 2021; Liu et al., 2025). We tested the effect of adipocyte differentiation factor PPARG, hematopoietic and endodermal organ differentiation factor HHEX, and muscle differentiation factor MYOD1 on expression of VISTA in A375, HCT116 or SK-HEP1 cells. There was a significant upregulation of VISTA expression in A375 and SK-HEP1 cells in response to forced expression of PPARG, upregulation of VISTA in A375 and HCT116 cells in response to forced expression of HHEX, and upregulation of VISTA in A375, HCT116 and SK-HEP1 cells in response to forced expression of MYOD1 (Fig. 4A). These data showed a general tendency that VISTA expression can be induced by non-neural pro-differentiation factors. Then we tested how oncoproteins SOX2 and KRAS affect expression of VISTA. SOX2 is a typical protein promoting neural stemness and pluripotency, and KRAS is primarily expressed in embryonic neural cells (Zhang et al., 2017). VISTA expression was suppressed by forced expression of SOX2 in A375, HCT116 and SK-HEP1 cells. Likewise, forced expression of both wild type KRAS and its oncogenic mutant KRAS(G12D) led to decreased expression of VISTA in cancer cells (Fig. 4B), showing that oncogenic proteins suppress VISTA expression. Regulatory effects of non-neural pro-differentiation factors and oncoproteins, which are embryonic neural or neural stemness factors, on VISTA expression explains why VISTA is detected primarily in differentiated cells in xenograft tumors.

**Fig. 4.**
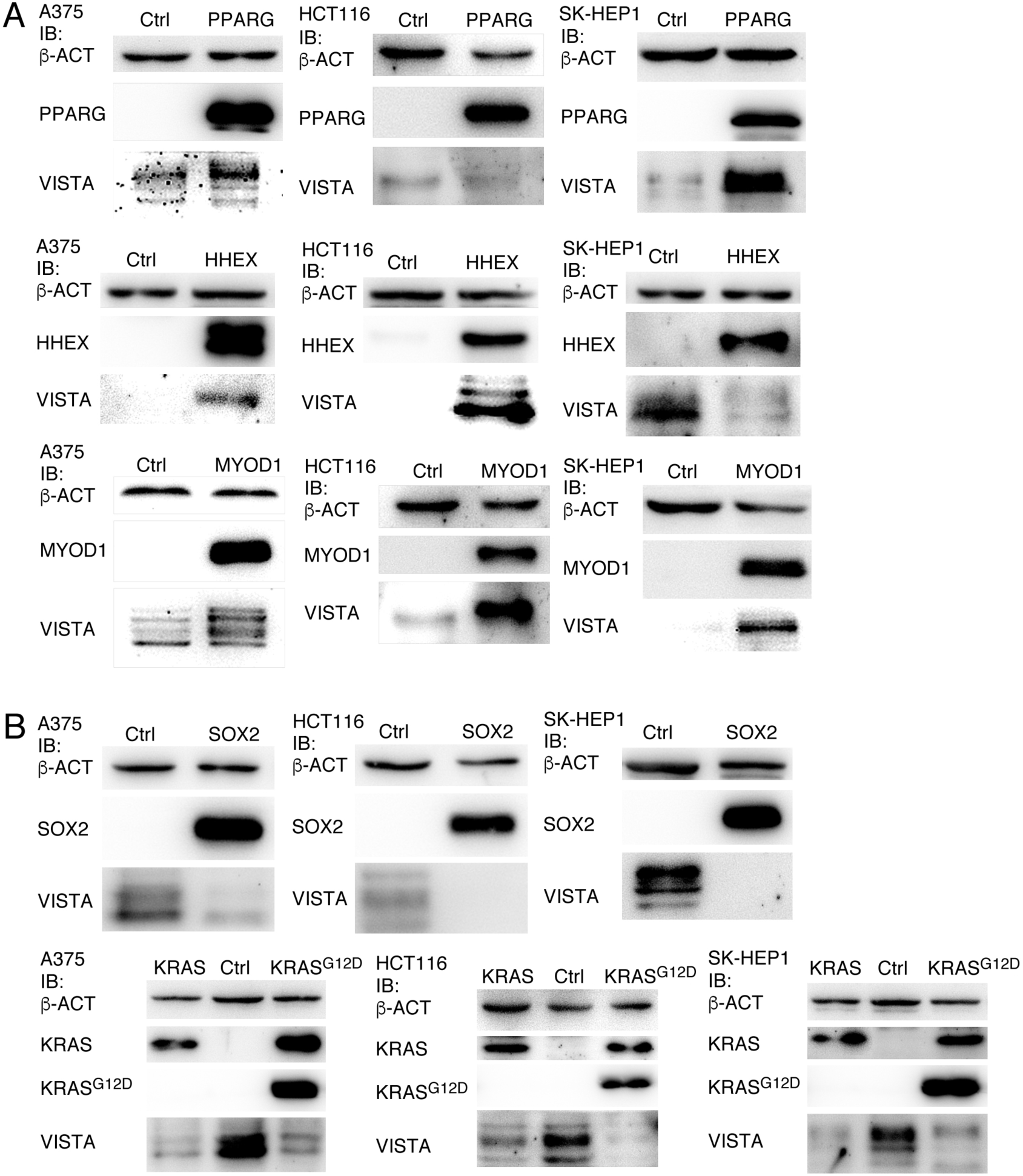
Differential regulation of VISTA by non-neural differentiation factors and oncoproteins. (A) Regulatory effect on VISTA of forced expression of non-neural differentiation factors, as revealed by IB. (B) Regulation of VISTA by forced expression of proteins promoting cancer, as revealed by IB. In each experiment in (A) or (B), control cells (Ctrl) and cells with forced expression of a protein were collected and subjected to IB. β-ACT was used as a loading control.

### VISTA regulates cell morphology

Manipulation of VISTA expression in cancer cells leading to change in cell morphology (Fig. 2) prompted us to investigate further whether VISTA is a regulator of cell morphology. C2C12 myoblast cells differentiated into muscle cells and formed long myotubes when cultured in medium containing low concentration of horse serum. When Vista expression was blocked, C2C12 cells still formed myotubes, but they appeared shorter and thicker than those formed by control cells (Fig. 5A, B). Staining of the differentiated muscle cell markers Myoglobin and Mhc (Myosin heavy chain) revealed that they were strongly expressed in myotubes formed by either control or Vista knockdown cells (Fig. 5A, B), suggesting that Vista inhibition does not affect differentiation of myoblasts, but affects the morphology of differentiated cells.

**Fig. 5.**
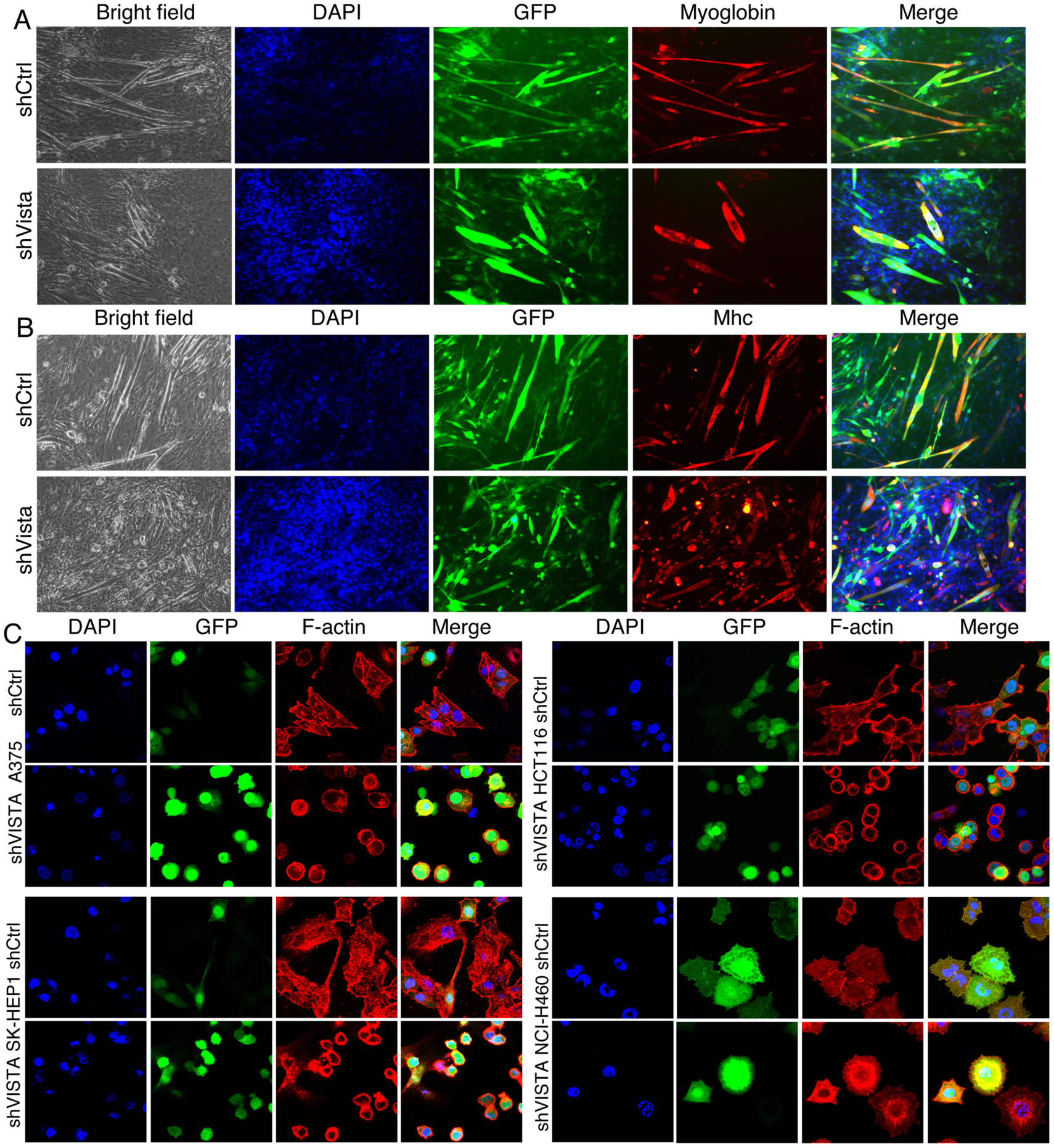
Effect of Vista/VISTA inhibition on cell morphology and actin cytoskeleton. (A) Morphological change in myotubes formed by C2C12 cells in response to knockdown of Vista, and expression of muscle cell markers detected with immunofluorescence (IF) in myotubes. (B) IF detection of F-actin in control cells (shCtrl) and cells with knockdown of VISTA (shVISTA).

Because cytoskeleton is crucial for maintaining cell morphology, we detected whether cytoskeleton was changed when VISTA was blocked. In control A375 cells, F-actin staining revealed clear distribution of actin filaments in cytoplasm and the cortical cytoskeleton. By contrast, actin filaments in cytoplasm of A375 cells with VISTA knockdown showed biased distribution toward one side of the cells. Moreover, staining of F-actin also clearly indicated a spherical morphology of these cells (Fig. 5C). Actin filaments also showed ubiquitous distribution in cytoplasm, cortical cytoskeleton and cell protrusions in control HCT116 cells, but they mainly distributed in cortical cytoskeleton in VISTA knockdown cells, which showed a spherical morphology and reduced cell protrusions (Fig. 5C). Ubiquitous distribution of actin filaments was more pronounced in control SK-HEP1 cells. Inhibition of VISTA led to a similar pattern of distribution of actin filaments as in A375 and HCT116 cells (Fig. 5C). No significant morphological change was not observed in NCI-H460 cells when VISTA was inhibited. However, alteration in distribution of actin filaments was still observed, as revealed by even distribution of actin filaments in control cells but intensified distribution in cytoplasm in knockdown cells (Fig. 5C). Change in cytoskeleton of cells in response to VISTA inhibition clearly indicates a role of VISTA in regulation of cell morphology.

### VISTA interacts with WASP2 in regulating cell morphology

To understand how VISTA affects cell morphology, RNAseq was carried out to analyze whether genes involved in regulating cytoskeleton could be changed in cells in response to VISTA inhibition. It was revealed that knockdown of VISTA led to downregulation of *WASF2* in A375, HCT116, SK-HEP1 or NCI-H460 cells (Fig. 6A), an effect that was confirmed by qRT-PCR (Fig. 6B). WASF2, also known as WAVE2 or WASP2, is a member of the Wiskott-Aldrich Syndrome Protein (WASP) family and functions primarily as a key regulator of the actin cytoskeleton (Soderling and Scott, 2006; Takenawa and Miki, 2001). In either HEK293T or HCT116 cells, forced expression of myc-tagged VISTA (VISTA-MT) led to enhanced expression of WASF2 (Fig. 7A). It can be noted that extranuclear distribution of VISTA and WASF2 is prominent (Fig. 7A). Such a result let us investigate whether VISTA and WASF2 interacts at protein level. Protein co-immunoprecipitation (co-IP) assays showed that VISTA-MT was able to precipitate endogenous WASF2, or vice versa, a myc-tagged WASF2 (WASF2-MT) was able to precipitate endogenous VISTA (Fig. 7B). To confirm further the involvement of VISTA in regulating actin cytoskeleton, thereby regulating cell morphology, via regulation of WASF2, we explored whether WASF2 could rescue morphological change in cells with inhibition of VISTA. In A375 cells, lentiviral particles for shVISTA were co-transfected with lentiviral particles for WASF2 at different MOI. shVISTA alone caused a spherical morphology in A375 cells. Co-transfection of WASF2 at 1 or 4 MOI didn’t rescue morphological change. When the dose of WASF2 virus particles was increased to 7 or 10 MOI, rescue effect on cell morphology became significant (Fig. 7C). Taken together, these results demonstrate that VISTA regulates actin cytoskeleton, thereby regulating cell morphology, via regulating and interaction with WASF2.

**Fig. 6.**
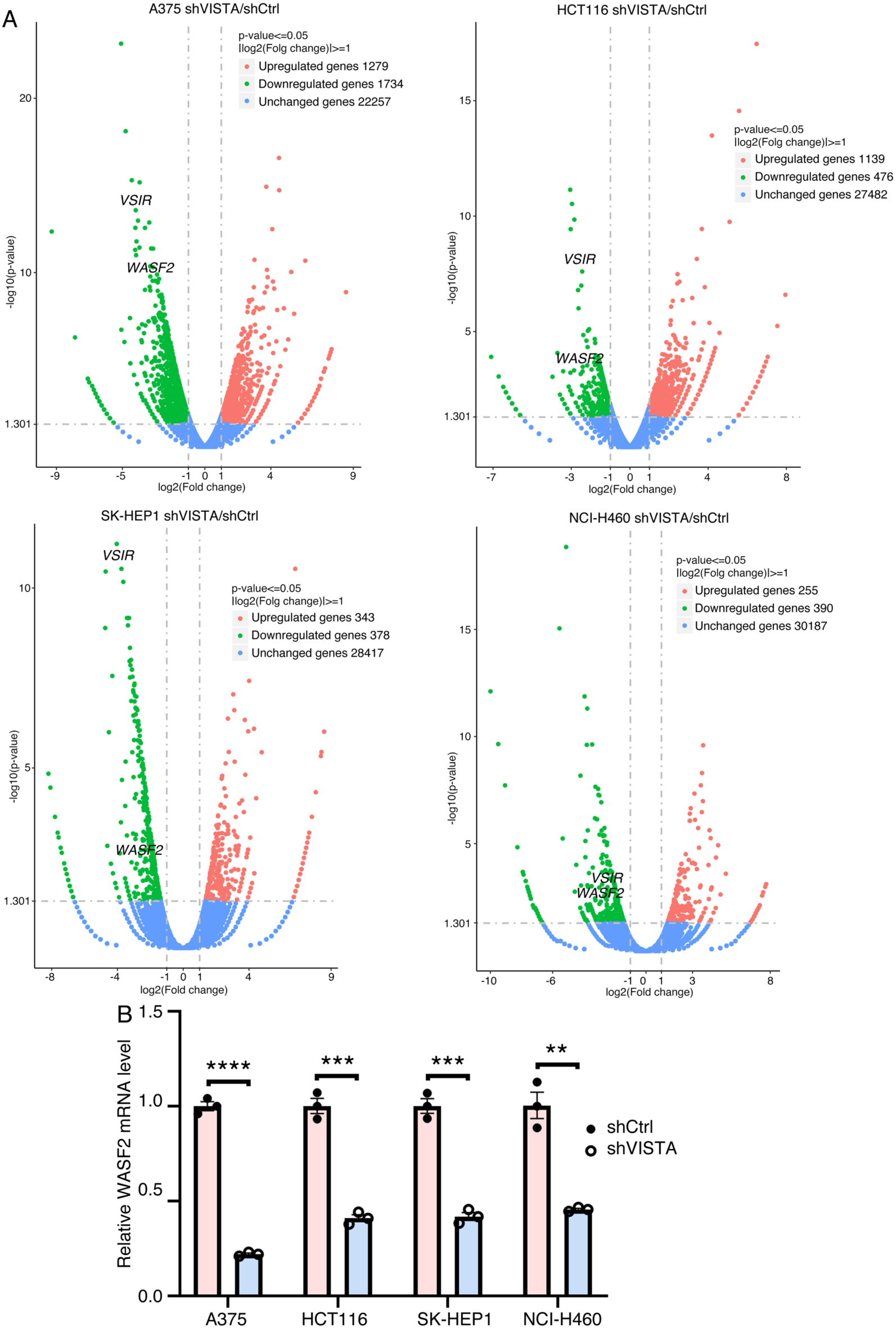
Identification of *WASF2* as a VISTA regulated gene. (A) Volcano plots generated from transcriptomic profiling showing downregulation of *WASF2* in all four cells with knockdown of VISTA/VSIR. (B) Confirmation of *WASF2* transcription in all four cells as in (A) in response to VISTA/VSIR knockdown with RT-qPCR. Significance in change of transcription levels was calculated based on experiments in triplicate using unpaired Student’s *t*-test. Data are shown as mean ± SEM. **p < 0.01, ***p < 0.001, ****p < 0.0001.

**Fig. 7.**
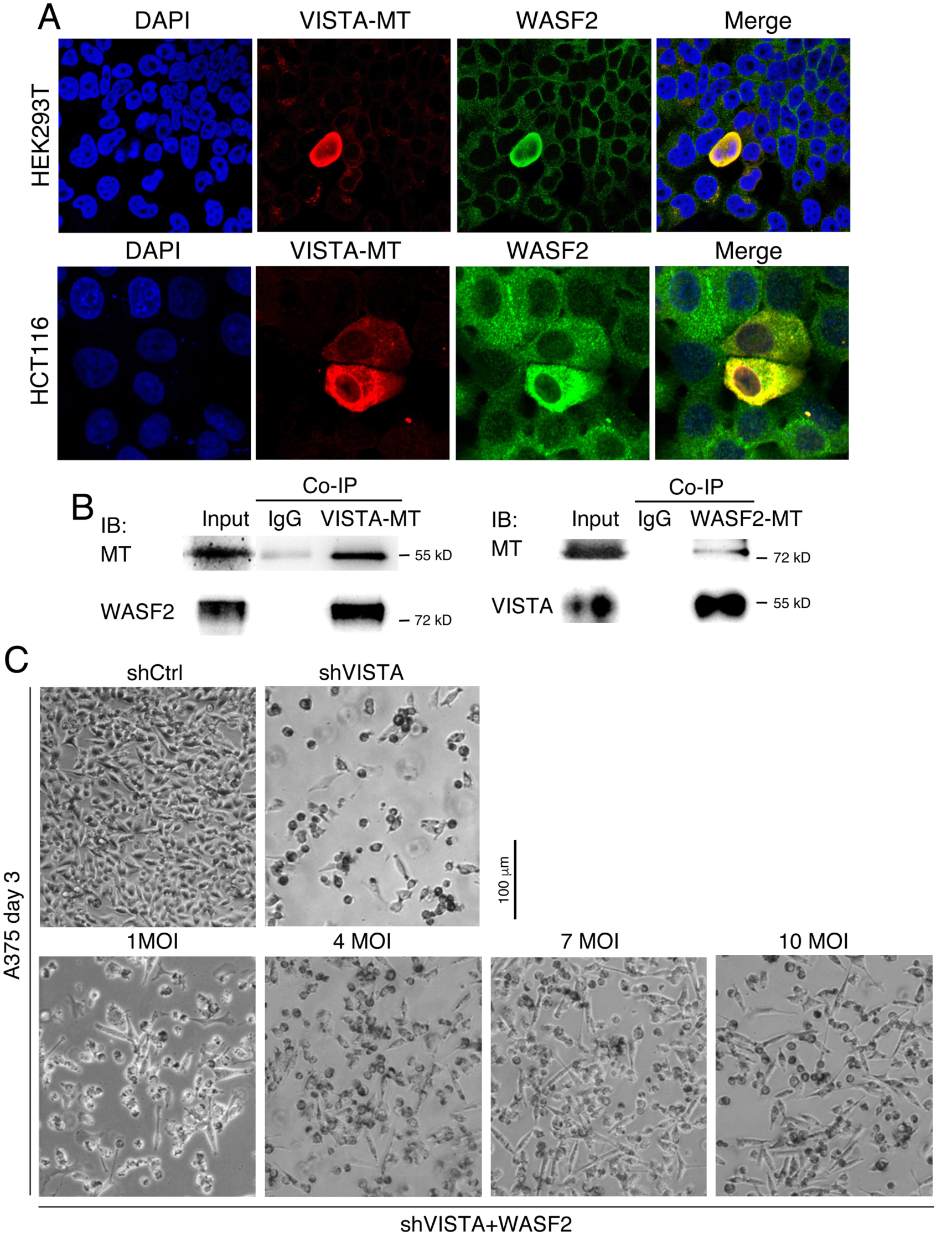
Regulation and interaction between VISTA and WASF2. (A) Overexpression of VISTA led to upregulation of WASF2 in both HEK293T and HCT116 cells, as revealed by IF assays. (B) Protein co-IP assays showing VISTA was able to precipitate WASF2, or vice versa. (C) Rescue of the morphological alteration in A375 cells resulting from VISTA knockdown by overexpression of WASF2 via transfection of virus at different MOIs as indicated.

## Discussion

### Overexpression of a gene in tumor might be the consequence of cancer cell differentiation and represent a less or non-malignant state

VISTA was considered as a therapeutic target mainly due to its function in regulating anti-tumor immunity and its overexpression in some types of cancer. However, a protein usually plays multiple roles during either embryogenesis or normal physiological/pathological processes. By analyzing expression patterns of SETDB1, which represents undifferentiated, malignant and immunoevasive state of cancer cells, and VISTA in xenograft tumors, higher levels of VISTA expression were observed only in cells without SETDB1 expression. This means that VISTA expression represents a more differentiated and less malignant state of cells, as compared with cells with SETDB1 expression. The core property of cancer cells is neural stemness, which confers cancer cells with pluripotency, and differentiation of cancer cells follows the principle of differentiation of embryonic pluripotent cells. Therefore, major non-neural lineage differentiation factors, e.g., HHEX, PPARG, MYOD1, could induce a higher level of expression of VISTA in cells, in agreement with expression of VISTA in a wide range of normal tissues (Martin et al., 2023). By contrast, VISTA expression is suppressed by oncoproteins SOX2 and KRAS, both are enriched in embryonic neural cells and play essential roles in regulating neural stemness (Bender et al., 2015). It can be concluded that VISTA expression tends to be upregulated by non-neural pro-differentiation factors but suppressed by oncoproteins, reinforcing that VISTA expression represents a differentiated and less malignant state in tumors. Further validation is from the effect of manipulated VISTA expression in cancer cells. Both forced expression and blockade of VISTA can cause morphological changes in cells but without significant differentiation effect and change in tumorigenicity. Interestingly, overexpression of Vista does not lead to change in tumor formation in syngeneic mouse models, suggesting that increase in VISTA level might not alter the response of cancer cells to anti-tumor immunity. Nevertheless, this evaluation might need more detailed experiments. These results suggest that if VISTA is chosen as a target of cancer therapy, only subpopulations of cancer cells with less malignancy and tumorigenicity are targeted but leaving cancer cells with higher malignancy and tumorigenicity untouched. It is well documented that cancer cell plasticity and intratumoral phenotypic heterogeneity increase with cancer progression (Meacham and Morrison, 2013; Dagogo-Jack and Shaw, 2018). Such a tendency is actually the reflection that neural stemness, hence the pluripotent differentiation potential, of cancer cells enhances progressively, thereby contributing to higher level of phenotypic heterogeneity during disease progression (Zhang et al., 2022; Cao, 2026). Therefore, some genes that are usually expressed neuronal and non-neural lineage differentiation are detected at high levels in tumor cells in high-grade disease, and their expression is also correlated with poor prognosis, e.g., the aforementioned TUBB3 and ITGB6. Choosing ITGB6 as a therapeutic target means that cells with least malignancy are to be wiped out but malignant ones still remain because ITGB6 is mainly expressed in muscle (Thisse and Thisse, 2004) and its expression represents muscle cell-like differentiation in tumors. So far, clinical trials of targeting ITGB6 in cancer showed only modest or even no significant activity in promoting patient survival (https://allsci.com/news/clinical-trials/pfizers-seagen-acquired-ib6-adc-fails-primary-survival-test-in-advanced-lung-cancer/). We propose that high expression of a gene and correlation between its expression and poor prognosis might not be a sufficient criterion for target selection. An additional factor should be whether expression of the gene of interest represents the most malignant cancer cell populations. This needs to understand the basic principles that most cancer promoting genes are embryonic neural/neural stemness genes but suppressor genes are mostly not, and neural stemness is the core property of cancer cells (Zhang et al., 2017; Cao, 2017; Xu et al., 2021; Cao, 2022).

### Immune checkpoints might play roles other than regulating anti-tumor immunity

Immune checkpoints were initially identified as regulators of anti-tumor immunity. Their role in cancer therapy has always been of particular interest to cancer researchers. Despite the initial success in immunotherapies with PD-1/PD-L1 and CTLA-4 blockade, the therapies have exhibited inefficiency in most cancer patients or resistance develops after therapy, or even cause an adverse effect of hyperprogression (Kamada et al., 2019; de Miguel and Calvo, 2020; Morad et al., 2021; Aliazis et al., 2025). It should be noted that it’s usual for a protein to play multiple roles and regulate different cellular functions or processes, and more importantly, cancer is not merely a disease of immunity. Besides the function in suppressing anti-tumor immunity, PD-1/PD-L1 were shown to function as tumor suppressors (Wang et al., 2020; Cao et al., 2021), making it complicated to evaluate the effect of inhibition of these checkpoints in cancer treatment. The present study showed that VISTA regulates cell morphology, as shown in cancer cells and muscle cell differentiation, via regulating actin cytoskeleton. At molecular level, the regulation is achieved through interaction with WASF2, a key regulator of cytoskeleton. Since interactions of surface molecules on adjacent cells, such as ligand/receptor interaction, is required for immune regulation, it’s reasonable that morphological change of cells would inevitably interfere with ligand/receptor interactions, leading to failure in signal transduction. VISTA blockade will probably fail to activate antitumor immunity because of its function in regulating cell morphology.

Previous focusing on cancer cells didn’t generate innovative new therapeutic strategies that are broadly beneficial across different cancers and significantly prolong overall survival (Swanton et al., 2024). This leads to the view that cancer cells themselves should not be the focus of the problem, and novel and more efficient therapeutic strategies should be developed based on the systemic complexity of cancer, especially the tumor microenvironment (TME) (Swanton et al., 2024). But contrary to what has been expected, novel strategies targeting components in the TME, such as inhibition of novel immune checkpoints, IDO1, TIGIT or LAG-3, have not achieved optimistic results, either. Given the endless complexity of TME and its regulatory molecular mechanisms, it raises a question how impractical it is to manipulate TME molecularly so that cancer can be more effectively suppressed. Logically, the central part of cancer should be cancer (tumorigenic) cells. When choosing a molecular target, whether the cells expressing the molecule are the (most) malignant ones should be considered. It should be kept in mind that expression of a gene could be the consequence of cancer (tumorigenic) cell differentiation during disease progression. Moreover, many ‘immune regulators’ play multiple functions in addition to regulation of immunity, and the underlying mechanisms cannot be simply understood in a linear way. Previous inefficiency in developing therapeutic strategies that achieve better benefit in different cancers might be the consequence of inadequate understanding of the basic property of cancer cells and the basic principles underlying tumorigenesis (Cao, 2017; 2022; 2023; 2026). Our identification of neural stemness as the core property of cancer cells and the basic rules underlying tumorigenesis suggest that neural stemness should be the therapeutic target (Yang et al., 2021; Cao, 2026).

## Methods

### Cell culture

A375, HEK293T, HCT116, SK-HEP1, and B16F10 cells were cultured in Dulbecco’s modified eagle medium (DMEM. Thermo Fisher Scientific, #11965092), NCI-H460, BxPC3, CT26, and 4T1 were cultured in RPMI-1640 medium (Thermo Fisher Scientific, #11875093), K562 was cultured in Iscove’s Modified Dulbecco’s Medium (IMDM. Thermo Fisher Scientific, #12440053), and NE-4C cells were in MEM (Gibco, #11090073) added with 1% MEM non-essential amino acids (Thermo Fisher Scientific, #11140050) and 1% Glutamax (Gibco, #35050061). All media were supplemented with 10% fetal bovine serum (FBS. Gibco, #10099141) and 50 U/ml penicillin/50 µg/ml streptomycin. All cells were cultured at 37°C with 5% CO_2_.

HEK293T (Cat. No.: #SCSP-502), HCT116 (Cat. No.: #TCHu99), SK-HEP1 (Cat. No.: #TCHu109), BxPC3 (Cat. No.: #TCHu12), K562 (Cat. No.: #TCHu191), NCI-H460 (Cat. No.: #TCHu205), B16F10 (Cat. No.: #SCSP-5233), CT26 (Cat. No.: #TCM37), and 4T1 (Cat. No.: #TCM32) were purchased from the Cellbank of Chinese Academy of Sciences (Shanghai, China). Cancer cell lines were authenticated with short tandem repeat profiling. Cells with fewer than eight passages were used for experiments.

### Plasmid construction, virus packaging, and cell transduction

Short-hairpin RNAs (shRNAs) used for knockdown of human VISTA or mouse Vista were designed with an online shRNA designer (https://portals.broadinstitute.org/gpp/public/seq/search). shRNAs were subcloned to the lentiviral vector pLKO.1, designated as shVISTA and shVista, respectively. Sequence of shVISTA is: GCACGATGTGACCTTCTACAA; shVista is: GGGAACCCTGCTCCTTGCTATT.

For stable expression of VISTA/Vista, SOX2, KRAS, KRAS(G12D), or WASF2, the coding regions of these genes or KRAS with the G12D mutation were subcloned to lentiviral vectors pLVX-IRES-Puro or pLVX-IRES-ZsGreen. Lentiviral vectors for stable expression of MYOD1, PPARG or HHEX were generated previously (Yang et al., 2021; Liu et al., 2025). For transient expression of VISTA or WASF2 in protein co-immunoprecipitation assays, the coding region of VISTA or WASF2 was subcloned to pCS2+6×MTmcs vector that contains six repeats of myc-tags (Zhang et al., 2022), and designated as VISTA-MT or WASF2-MT.

For lentivirus production, HEK293T cells were co-transfected with packaging plasmids and shRNA or expression constructs in the presence of polyethylenimine (PEI). 48 hours after transfection, lentiviral supernatant was filtered through 0.45 μm filters and centrifuged at 4°C to concentrate viral particles. For cell transfection, polybrene at a final concentration of 10 µg/ml was used. Cells were selected with puromycin 48 hours after viral transfection when a puromycin selection vector was used, and were cultured further for observation of phenotypic change or for additional assays. As a control, virus production with an empty vector and cell transfection were performed in parallel. In rescue assays, A375 cells were first transfected with viral particles for shVISTA. One day later, viral particles for expression of mouse Vista or human WASF2 were transfected at different multiplicity of infection (MOI), as indicated in text.

For transient overexpression assays, HEK293T cells or HCT116 cells were transfected with VISTA-MT or WASF2-MT plasmid or a vector plasmid (as a control) using PEI when cells grew to 70 to 80% confluency. Cells were collected for further assays 48 hours later after plasmid transfection.

### Cancer cell transplantation in mice

Mouse use was approved by the Institutional Animal Care and Use Committee (IACUC) at the Model Animal Research Center, Medical School of Nanjing University, and experiments with mice were performed in accordance with the guidelines of IACUC. Immunodeficient nude Foxn1nu male mice, C57BL/6J, and BALB/c mice at 5-6 weeks were purchased from the National Resource Center for Mutant Mice (Nanjing, China) and maintained in a specific-pathogen-free facility. Control or treated cancer cells were suspended in 100 µl of PBS and injected subcutaneously into the dorsal flank of a mouse. Tumor formation was observed and tumor size was measured periodically. After sacrifice of mice, tumors were excised. Tumor volume was calculated with the formula: length×width2/2. Significance of difference in tumor volume between control and treated groups was calculated with two-way ANOVA-Bonferroni/Dunn test. Control or treated HCT116 cells were injected at a dose of 3×10^6^ cells per mouse, SK-HEP1 at a dose of 5×10^6^, K562 at a dose of 5×10^6^, A375 at a dose of 3×10^6^, NCI-H460 at a dose of 3×10^6^, BxPC3 at a dose of 1×10^7^, NE-4C at a dose of 1×10^6^, 4T1 at a dose of 5×10^5^, B16F10 at 5×10^5^, and CT26 injected at a dose of 5×10^5^ per mouse.

### Immunohistochemistry (IHC)

IHC detection of protein expression in paraffin sections of tumors was performed using conventional method. Briefly, paraffin in sections was removed by washing in xylene for 10 min first and then 5 min. Sections were rehydrated with sequential washes with decreasing gradient of in 100%, 95%, 85%, 70%, 50% ethanol, and dH_2_O for 5 min each. For antigen retrieval, sections were treated with 0.01 M sodium citrate solution at 95-100°C for 20 min, then rinsed with PBS at room temperature. Sections were treated with 3% H_2_O_2_ in methanol for 15 min to block endogenous peroxidase activity. After washing slides with PBS three times, sections were blocked with 5% BSA in PBS for 1 hour. Afterwards, primary antibody was added and incubated at 4°C overnight, followed by washing with PBS. Subsequently, biotin-conjugated goat anti-rabbit secondary antibody (Sangon Biotech, #D110066. 1:500) was added to sections, which were incubated at room temperature for 1 hour. Signals were revealed with addition of a DAB substrate (Sangon Biotech, #E670033). Cell nuclei were counterstained with hematoxylin. Primary antibodies were: SETDB1 (Cell Signaling Technology, #2196. 1:200), VISTA (Cell Signaling Technology, #54979. 1:1,000)

### Immunoblotting (IB)

IB was performed using conventional method. Briefly, cells were trypsinized and then lysed with RIPA buffer. Lysates were centrifuged at 4°C to remove cell debris. Afterwards, 5×loading buffer was added to the supernatant, followed by denaturation at 98°C. Protein samples were then subjected to SDS-PAGE, transferred to PVDF membrane, which was blocked with blocking buffer (5% non-fat milk in TBST). Primary antibody was then added to the membrane that was submerged in blocking buffer, incubated at 4°C overnight. After washing the membrane three times with TBST, secondary antibody was added, incubated for 2 hours at room temperature. Membrane was washed three times, and protein bands were revealed with a Western blotting substrate (Tanon, #180-501). The primary antibodies were: β-ACT (Abclonal, #AC004. 1:10,000), VISTA (Cell Signaling Technology, #54979. 1:1,000), PCNA (Cell Signaling Technology, #13110. 1:2,000), PPARG (Cell Signaling Technology, #2435. 1:1,000), SETDB1 (Cell Signaling Technology, #2196. 1:1,000), EZH2 (Cell Signaling Technology, #5246. 1:2,000), DNMT1 (ABclonal, A16729, 1:1,000), MYOD1 (Novus Biologicals, NBP1-54153. 1:1,000), MSI1 (ABclonal, A9122. 1:1,000), C-MYC (Abcam, #ab42072. 1:1,000), HHEX (R&D, #MAB83771. 1:2,000), KRAS (Abcam, #ab172949. 1:2,000), KRAS(G12D) (Abcam, #ab221163. 1:2,000), SOX2 (Abcam, #ab92494. 1:1,000), WASF2 (Cell Signaling Technology, #3659. 1:2,000), Myc-tag (Abclonal, #AE010. 1:1,000)

### F-actin staining

Control or treated cells were grown to 50% confluence on slides that were pre-coated with poly-D-lysine. Cells were washed twice with PBS, and fixed with 4% formaldehyde, followed by washing again with PBS for three times. Afterwards, cells were permeabilized with 0.5% Triton X-100 in PBS for 5 min, and washed twice with 10 min each. Cells were stained with TRITC-Phalloidin (Yeasen, #40734ES) at 20 nM for 30 min under dark, washed two time with 0.5% Triton X-100, and three times with PBS. Cell nuclei were counterstained with DAPI. After mounting the slides, cells were observed and images were scanned under laser con-focal microscope (Leica).

### Myoblast differentiation

Control C2C12 myoblast cells and C2C12 with Vista knockdown were cultured in DMEM containing 2% horse serum (Sangon Biotech, #E510006). Differentiation was manifested by myotube formation.

### Immunofluorescence (IF)

Cells were fixed with 4% PFA for 15 min, washed twice with PBS, and permeabilized with 0.1% Triton X-100 at room temperature for 10 min, and followed again by washing twice with PBS. After blocking cells with 0.2% gelatin in PBS for 30 min, primary antibody was added and incubated overnight at 4°C. Primary antibody was removed by washing cells three times with PBS. Afterwards, secondary antibody was added and incubated at room temperature for two hours. Cell nuclei were counterstained with DAPI. Primary antibodies were: Myoglobin (Cell Signaling Biotechnology, #25919. 1:200), Mhc (R&D systems, #MAB4470. 1:200), WASF2 (Cell Signaling Technology, #3659. 1:200), and Myc-tag (Abclonal, #AE010. 1:200)

### Protein co-immunoprecipitation (Co-IP)

Co-IP was performed essentially as described (Chen et al., 2021). Cells were collected and washed twice with ice-cold PBS, and lysed with NP-40 lysis buffer on ice for 20 min. Cell lysates were centrifuged for 20 min at 12,000 rpm. Supernatants were collected and immunoprecipitation was performed using an antibody against Myc-tag (Abclonal, #AE010) that was linked to protein A/G sepharose beads. Immunoprecipitation with the antibody against IgG was performed in parallel as negative controls. After incubation at 4°C overnight, beads were collected by centrifugation and washed with TBST buffer. Immunoprecipitates were eluted by incubating the beads in 1 × loading buffer at 95°C for 8 min, and then subjected to SDS-PAGE and Western blotting.

### Reverse transcription-quantitative polymerase chain reaction (RT-qPCR)

Total RNA was prepared from cells using TRIzol reagent. Reverse transcription of complementary DNA (cDNA) was carried out using the HiScript II 1st Strand cDNA Synthesis Kit (+gDNA wiper) (Vazyme, #R212-01/02), which contains reagent for removing genomic DNA. qPCR was performed on a LightCycler96 System (Roche) using following parameters: one cycle of predenaturation at 95°C for 1 min, followed by 40 cycles of denaturation at 95°C for 10 s, and annealing and extension at 60°C for 30 s. Afterwards, one additional cycle for melting curve was performed. In each experiment, transcription of *β-ACT* was included as an internal control. Significance in difference of transcription level was calculated based on experiments in triplicate using unpaired Student’s *t*-test. Results are presented as histograms with relative units of transcription level. Primers were:

*β-ACT* (Forward): AGAAAATCTGGCACCACACC

*β-ACT* (Reverse): TAGCACAGCCTGGATAGCAA

*WASF2* (Forward): TGCAAGGAATCAACACCCGA

*WASF2* (Reverse): TGCCTCTTTTCCATCGTCCC

*VISTA* (Forward): TGTGGTGTACCCATCCTCCT

*VISTA* (Reverse): TTGAATGTTGCTGTCCATCC

### RNA sequencing (RNAseq)

In order to analyze the genes that are potentially regulated by VISTA, RNAseq was performed with A375, HCT116, SK-HEP1 and NCI-H460 cells. Control cells and cells with VISTA knockdown were collected, and total RNA was extracted with TRIzol. Quantity and quality of RNA were evaluated with RNA Nano 6000 Assay Kit of the Bioanalyzer 2100 system (Agilent Technologies, CA, USA). mRNA was purified, and first strand cDNA was synthesized from fragmented mRNA. After removal of RNA, second strand cDNA was synthesized. After qualification, cDNA library was subjected to sequencing by Illumina NovaSeq 6000. Image data measured by the sequencer were converted into sequence by CASAVA base calling. Raw data were firstly processed through in-house perl scripts, and mapped against human reference genomes using Hisat2 (v2.0.5) software, and mapped reads were assembled with StringTie (v1.3.3b). FPKM of each gene was calculated based on the length of the gene and reads count mapped to this gene. Differential expression was analyzed with the edgeR package (3.24.3). P values were adjusted with the Benjamini-Hochberg method. Padj≤0.05 and |log2(fold change)| ≥1 were set as the threshold for significantly differential expression.

RNA extraction, library construction, sequencing, signal processing and data analyses, including volcano plots, were performed by Novogene Co., Ltd. (Beijing, China). RNAseq data were deposited to Gene Expression Omnibus (GEO) under accession numbers: GSE344143, GSE344144, GSE344145, and GSE344146.

## Author contributions

CW: Formal Analysis, Investigation, Visualization, Writing; YL: Investigation; JL: Investigation; YC: Conceptualization, Formal Analysis, Funding acquisition, Resources, Visualization, Writing.

## Acknowledgments

We would like to thank the voluntary assistance of Haihua Ma, and thank all who supported and encouraged our research in different ways on the topic “Neural stemness as the core property of cancer cell”.

